# Small gene networks can delineate immune cell states and characterize immunotherapy response in melanoma

**DOI:** 10.1101/2022.07.11.498823

**Authors:** Donagh Egan, Martina Kreileder, Myriam Nabhan, Luis F. Iglesias-Martinez, Simon Dovedi, Viia Valge-Archer, Amit Grover, Robert Wilkinson, Tim Slidel, Claus Bendtsen, Ian Barrett, Donal Brennan, Walter Kolch, Vadim Zhernovkov

**Affiliations:** Precision Oncology Ireland, Systems Biology Ireland, School of Medicine, University College Dublin, Belfield, Republic of Ireland; Catherine McAuley Research Centre Mater Misericoridae University Hospital, Dublin, Republic of Ireland; AstraZeneca, Cambridge Biomedical Campus, Cambridge, United Kingdom; Conway Institute of Biomolecular & Biomedical Research, University College Dublin, Belfield, Dublin 4, Ireland

**Keywords:** Immune Checkpoint Inhibitors, Gene Networks, Single Cell RNAseq, Bioinformatics, Computational Modelling, Melanoma

## Abstract

**Background:** Single-cell sequencing studies have elucidated some of the underlying mechanisms responsible for immune checkpoint inhibitor (ICI) response, but are difficult to implement as a general strategy or in a clinical diagnostic setting. In contrast, bulk RNAseq is now routine for both research and clinical applications. Therefore, our analysis extracts small transcription factor-directed co-expression networks (regulons) from single-cell RNA-seq data and uses them to deconvolute immune functional states from bulk RNA-seq data to characterize patient responses.

**Methods:** Regulons were inferred in pre-treatment CD45+ cells from metastatic melanoma samples (n=19) treated with first-line ICI therapy (discovery dataset). A logistic regression-based classifier identified immune cell states associated with response, which were characterized according to differentially active, cell-state specific regulons. The complexity of these regulons was reduced and scored in bulk RNAseq melanoma samples from four independent studies (n=209, validation dataset). Patients were clustered according to their regulon scores, and the associations between cluster assignment, response, and survival were determined. Intercellular communication analysis of cell states was performed, and the resulting effector genes were analyzed by trajectory inference.

**Results:** Regulons preserved the information of gene expression data and accurately delineated immune cell phenotypes, despite reducing dimensionality by > 100-fold. Four cell states, termed exhausted T cells, monocyte lineage cells, memory T cells, and B cells, were associated with therapeutic responses in the discovery dataset. The cell states were characterized by seven differentially active and specific regulons that showed low specificity in non-immune cells. Four clusters with significantly different response outcomes (P <0.001) were identified in the bulk RNAseq validation cohort. An intercellular link between exhausted T cells and monocyte lineage cells was established, whereby their cell numbers were correlated, and exhausted T cells predicted prognosis as a function of monocyte lineage cell number. Analysis of ligand – receptor expression suggested that monocyte lineage cells drive exhausted T cells into terminal exhaustion through programs that regulate antigen presentation, chronic inflammation, and negative co-stimulation.

**Conclusions:** Regulon-based characterization of cell states provides robust and functionally informative markers that can deconvolve bulk RNA-seq data to identify ICI responders.

## INTRODUCTION

Despite the great success of immune checkpoint inhibitors (ICIs), only 12% of patients across all cancer indications are estimated to respond to therapy.^1^ The tumor microenvironment (TME) plays an important role in response outcomes to ICIs, exerting both immunostimulatory and immunosuppressive effects.^2^ ICI therapies aim to counteract the dysfunctional effects of the TME on T cells.^3^ Understanding which immune cell states are amenable to functional revival, which cell states are refractory, and how susceptible and resistant cell states dynamically interact, could provide new opportunities for disease management and therapeutic intervention.

Several studies have analyzed the correlation between immune cell states and ICI response. For instance, effector and partially exhausted CD8+ T cells incite tumor control in response to ICIs.^4^ However, when differentiated into a terminally exhausted state, due to continued antigen stimulation, they fail to control tumor progression.^5^ Regulatory T cells (Tregs) attenuate the activity of CD8+ T cells to maintain self-tolerance. Preclinical studies have demonstrated that anti-CTLA-4 monoclonal antibodies may improve responses, at least in part, through modulation of Treg-mediated suppression of effector CD8+ T cells.^6^ Tumor-associated macrophages (TAMs) represent a spectrum of phenotypes with M1 and M2 macrophages at either end. Inhibiting the activity of M2-like TAMs and redirecting their polarization towards the M1 phenotype can enhance response to ICIs.^7,8^ An outstanding question from such studies is whether tumor-infiltrating immune cell states, their transcriptional landscape, and co-occurrence patterns can effectively characterize patient response to ICIs.

Single-cell genomics at large-scale has emerged as a powerful technology for obtaining high-resolution transcriptomic data of tumor cellular ecosystems from primary tumor samples. However, constrained by prohibitive costs, single-cell RNA-seq (scRNA-seq) cohorts are often modestly sized, making it difficult to validate biomarkers for predicting response to drug treatments.

Here, we leveraged the resolution offered by scRNA-seq to identify immune cell states associated with response to ICIs, and the larger sample sizes of bulk RNA-seq datasets for validation. We demonstrated that a signature of transcription factor (TF)-directed gene networks, characterizing cell states associated with response, can identify patients who are likely to receive clinical benefit from ICI treatment. We explored dependencies between these cell states, and established an intercellular link between monocyte-lineage cells (MLCs) and exhausted T cells, which affects patient prognosis. Overall, these findings support the importance of considering the multifaceted nature of immune cell states present in the TME, and not limiting treatment strategies to T cells.

## MATERIALS AND METHODS

### Patient Samples from Previously Published Cohorts

Pre-treatment samples with response information available in the form of RECIST criteria were selected from published studies.^9–14^ The overall survival and clinical response data, as originally reported, were used. A binary definition of response was implemented to standardize clinical response across studies: complete response (CR), partial response (PR), or stable disease (SD) for responders, and progressive disease (PD) for non-responders.

### Single-cell RNA-seq Data Preparation and Regulon Inference

For the melanoma dataset, log-normalized transcripts per million (TPM) scRNA-seq data were accessed through NCBI’s Gene Expression Omnibus (GEO) (GSE120575) and processed as described in.^9^ An additional filtering step of genes expressed in <1% of cells or with <500 counts was implemented. A TF-directed co-expression network was inferred from the filtered gene expression matrix of pre-treatment samples using *GRNBoost2* in (https://github.com/tmoerman/arboreto) *py*SCENIC. *RcisTarget* (v1.6.0) was then used for cis-regulatory motif analyses on the co-expression network, removing indirect targets from TF modules to create regulons. The resulting regulon activity scores were calculated across each cell in the filtered gene expression matrix using *AUCell* with default parameters (v1.8.0).^15^

The basal cell carcinoma dataset was accessed from GEO under the accession number (GSE123814). The raw unique molecular identifier counts (UMI) were log-normalized, and genes expressed in <1% of cells or with <500 counts were removed. The regulons identified in the melanoma dataset were scored using *AUCell* with default parameters.

### Cell Clustering

Clustering was performed independently on the gene expression and regulon activity feature space using the Seurat package (Version 4.0).^16^ The gene expression and regulon activity values were first scaled using the “ScaleData” function.^17^ Clusters were identified using shared nearest neighbor (SNN)-based clustering, with the first 15 principal components, *k* = 20, and resolution = 0.3 being provided as inputs. The same principal components were used to generate the UMAP projections; all other parameters were set to their default values. The cell annotations were retained.^9^

### Identification of Differentially Abundant Cell States

The DAseq algorithm was used to identify differentially abundant (DA) cell states between responders and non-responders.^18^ The regulon UMAP projection previously generated from Seurat was used as input to DAseq. DAseq uses a k-nearest neighbor algorithm to define the local neighborhood of a cell from which its differential abundance score is calculated. The range of k values was set to start at 50 and end at 500, with a step of 50. A permutation test was used to determine the threshold for which a cell was considered DA. All other DAseq parameters were set to default values.

### Characterizing Cell States of Response and Regulon Pruning

Differentially active regulons between DA cell states from DAseq were calculated using the Wilcoxon test through the Seurat package.^16^ To quantify the cell state specificity of a regulon, we adapted an entropy-based strategy (the Jensen-Shannon divergence) that was previously described.^19^ For each cell state, regulons with the highest cell-state specific and differential activity scores were identified as essential regulators

Regulon modules were identified based on the Connection Specificity Index (CSI)-^20^ a context-dependent measure for identifying specific associating partners. The evaluation of the CSI was described previously.^19^

To reduce regulon complexity, whilst maintaining a cell-type-specific effect, target genes for each selected regulon were pruned if they were not upregulated in the predominant immune cell phenotypes of its associated cell state. Marker genes for each immune cell phenotype (original cell clusters) were determined according to Sade-Feldman et al.^17^

### Validation Datasets

RNA sequencing data of pre-treatment melanoma biopsies were obtained from the following studies: 1) Van Allen et al., an advanced melanoma anti-CTLA-4 treated cohort;^11^ 2) Hugo et al., an advanced melanoma anti-PD-1 treated cohort;^13^ 3) Riaz et al., an advanced melanoma anti-PD-1 treated cohort;^12^ 4) Gide et al., an advanced melanoma anti-PD-1 or combined ICI treated cohort.^14^ This combination of studies was chosen as the distribution of treatments received modelled the scRNA-seq discovery cohort (Supplementary Table1). Response was defined in the same way as the discovery cohort.

For Hugo et al. and Riaz et al., raw FASTQ files were obtained from the Sequence Read Archive (SRP067938, SRP094781). The RASflow pipeline was used for trimming, alignment, and feature counting.^21^ For Gide et al. and Van Allen et al., raw FASTQ files were pre-processed as described,^22^ and the processed counts were accessed at (https://zenodo.org/record/4661265). Gene counts for each dataset were normalized using the standard DEseq2 protocol.^23^

### Regulon Scoring, Batch Effect Correction, and Patient Clustering

The finalized regulon gene sets were scored independently in each validation dataset using Single Sample Gene Set Enrichment Analysis (ssGSEA).^24^ Median centering and scaling were performed on each dataset, which were then combined. Hierarchical clustering of the patients according to their regulon scores was performed using the Pheatmap R package. Average linkage clustering was applied with a correlation distance metric. Associations between cluster assignment and patient response were assessed using the chi-squared test.

### Modelling intercellular communication

The R package nichenetr (NicheNet; v1.0.0) was used to infer cell communication between the MLC and exhausted T cell states identified by DAseq.^18,25^ The MLCs were defined as the “sender” cells, and the exhausted T cells were considered as the “receiver” population. Genes with mean expression > 2 log_2_(TPM) in either cell type were retained, and those expressed in exhausted T cells were used as “background genes”. The “gene set of interest” required by nichenetr was defined as the RUNX3 and PRDM1 regulons. The top five ligands were selected according to their regulatory potential score, and were considered to promote the expression programs of RUNX3 and PRDM1.

### Trajectory Inference

Cytotoxic lymphocytes, exhausted CD8 T cells, exhausted heat shock CD8 T cells, lymphocytes exhausted/cell cycle and memory T cells, as defined by Sade-Feldman et al. using gene expression markers,^9^ were included in trajectory inference. Principal component analysis was applied to the gene expression matrix for these cell types; principal components 1 and 2 were used as inputs to a Gaussian mixture model to identify pseudo-time-dependent cell clusters (Mclust package).^26^ Slingshot was used to fit a minimum spanning tree (MST) to these clusters and determine the approximate trajectory.^27^ Memory T cells were set as the starting cluster. This piecewise linear trajectory was smoothed using simultaneous principal curves to obtain the final trajectories and pseudo-time values.

The associations between pseudo-time and individual gene expression levels were described by fitting negative binomial general additive models via the tradeSeq package.^28^ These models use cubic splines as basis functions to express a gene’s average expression level as a function of pseudo-time.

### Statistical Analysis

#### Survival Curves

Kaplan-Meier curves were generated using the Survival R package (Version 3.2-13) and compared using the log-rank test.

#### Cox Regression Model

The mean patient ssGSEA activity of RUNX3 and PRDM1, as inferred previously, was determined using the Van Allen dataset.^11^ Patients were split into high- and low-MLC cohorts according to the median score for the MAFB regulon. A Cox regression model was used to estimate the hazard ratios of deaths associated with a patient’s mean activity of RUNX3 and PRDM1 while accounting for their membership in either the high or low MLC cohorts.

#### Multiple Linear Regression

The top 5 regulatory potential scoring MLC ligands according to nichenetr, and a CIBERSORT “Absolute Score”, were used as explanatory variables in a multiple linear regression modelling the mean activity of RUNX3 and PRDM1 in the Van Allen dataset. The MLC ligands were scored using ssGSEA,^24^ and the “Absolute Score” was determined at [CIBERSORTx](https://cibersortx.stanford.edu/jobs.php) (stanford.edu), using the LM22 Source GEP.^29^

## RESULTS

### Regulon Inference and Cell States of Response

The presence or absence of certain immune cells in the TME influences response to ICIs.^2,30^ Identifying and characterizing these immune cells can provide avenues for understanding and predicting response to ICIs. We analyzed scRNA-seq data derived from tumor resident immune cells of metastatic melanoma patient samples (n=19) collected prior to ICI treatment.^9^ ScRNA-seq data are inherently noisy due to sparse and shallow sequencing with missing values, which can compromise the clear, robust, and reproducible distinction of different cell types. To circumvent this problem, we inferred TF-directed gene networks (hereafter referred to as regulons) using the SCENIC workflow.^15^ Regulons act as a buffer against noisy expression fluctuations at the individual gene level, remaining stable despite numerous resampling and multiple sample batches.^31^ A regulon is denoted by the upstream TF, contains its direct downstream gene targets, and reflects the activity of the TF across each cell.

The cells were clustered using either gene expression or regulon activity. A comparison of both approaches revealed that information was not sacrificed for the dimensionality reduction introduced by regulons: expression variability remained between immune cell phenotypes (Fig 1A).

**Figure 1:**
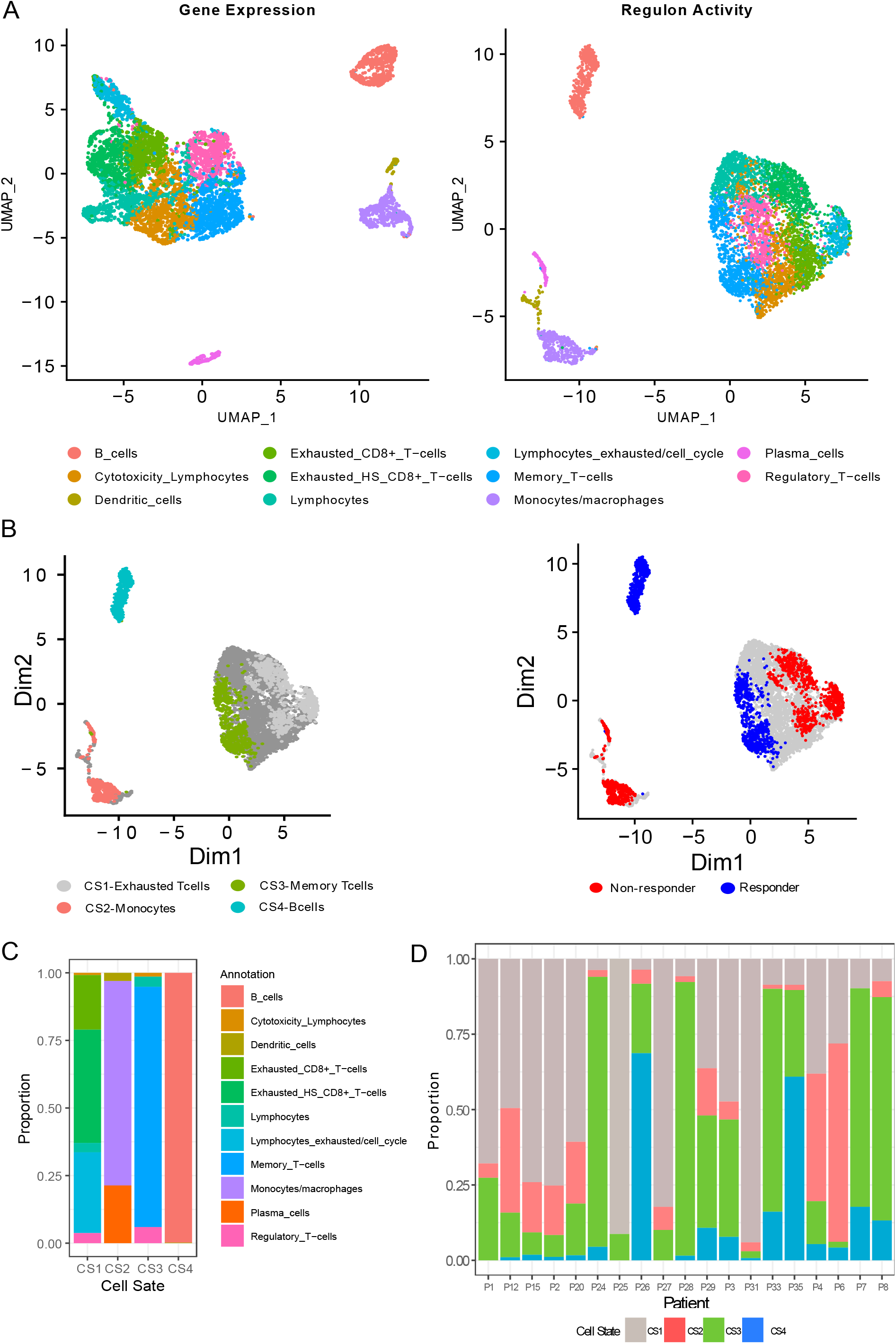
Clustering of cells and their association with response. **A)** Dimensionality reduction, clustering and UMAP projection of tumor-resident immune cells before ICI therapy for 19 melanoma patients. Left-sided figure: clustering according to gene expression profiles. Right-sided figure: clustering according to regulon activity profiles. **B)** UMAP embeddings of pre-treatment cells, annotated based on DAseq identified cell states (left panel), and their association with response (right panel). **C)** Distribution of immune cell phenotypes in the four DAseq identified cell states. **D)** The proportion of the DAseq identified cell states in each patient sample.

Leveraging the DAseq method,^18^ four distinct pre-treatment cell states were identified as DA between responders and non-responders (Fig 1B). RECIST criteria were used to define response, with stable disease, complete and partial response representing responders, and progressive disease representing non-responders. Each cell state was annotated according to its predominant immune cell phenotype (Fig 1C). Cell state 1 (CS1), termed Exhausted T cells, was associated with non-responders and contained three distinct exhaustion phenotypes: exhausted CD8 T cells, exhausted heat shock CD8 T cells, and lymphocytes exhausted/cell cycle. Annotated as monocyte lineage cells (MLCs), cell state 2 (CS2) contained predominately Monocytes/Macrophages and was associated with non-responders. Cell state 3 (CS3) was abundant in responders and consisted of memory T cells. B cells made up cell state 4 (CS4), whose presence was also associated with responders.

The proportion of cell states present in each patient sample was analyzed to ensure that the association of cell state with response was not driven by a limited number of patients. Importantly, each cell state contained cells from the majority of the patients (Fig 1D).

### Regulons Defining Cell States of Response

During immune cell differentiation from multipotent hematopoietic stem cells, lineage-restricted TFs are induced, and in turn establish identity of a specific cell type.^32^ We posit that regulons can characterize the transcriptional state of immune cells, providing a robust and functionally informative marker for deconvolving the TME. To identify regulons capable of characterizing the DAseq identified cell states, we employed two metrics: 1) differential regulon activity using a Wilcoxon test; 2) regulon specificity according to the Jensen-Shannon divergence.^33^ A differentially active regulon for a particular cell state has a statistically significant increase in median activity compared to all other cell states. Regulon specificity measures how distinct the distribution of regulon scores is for a specific cell state. Regulons scoring highly for both metrics in a cell state were selected.

The PRDM1 and RUNX3 regulons were significantly upregulated and specific to the exhausted T cell state (Fig 2A, 2B). Both TFs are known to modulate terminal differentiation of effector CD8 T cells, with RUNX3 being required for accessibility to PRDM1 transcription factor motifs.^34,35^ The LEF1 and FOXP1 TFs demonstrate enhanced expression in memory T cell phenotypes.^36^ Unsurprisingly, their associated regulons were highly specific and differentially active in the memory T cell state (Fig 2A, 2B). MAFB and PAX5 scored highly for both metrics in the MLCs and B-cell states, respectively (Fig 2A, 2B). MAFB is an inducer of monocyte differentiation,^37^ whilst PAX5 is a master regulator of B cell development.^38^ A proportion of the exhausted and memory T cell states (CS1 and CS3 clusters) identified by DAseq contained Tregs (Fig 1C). Tregs represent an important cell type in ICI response. To quantify this cell type in future deconvolution steps, SMAD1 was identified as being specific to and differentially active in Tregs (Fig 2A, 2B). TGF-β-dependent recruitment of SMAD has been identified as a key component in the generation of Tregs by participating in the “on-and-off” switch of FOXP3, and manipulation of SMAD’s activity is sufficient to alter the Treg differentiation program.^39^

**Figure 2:**
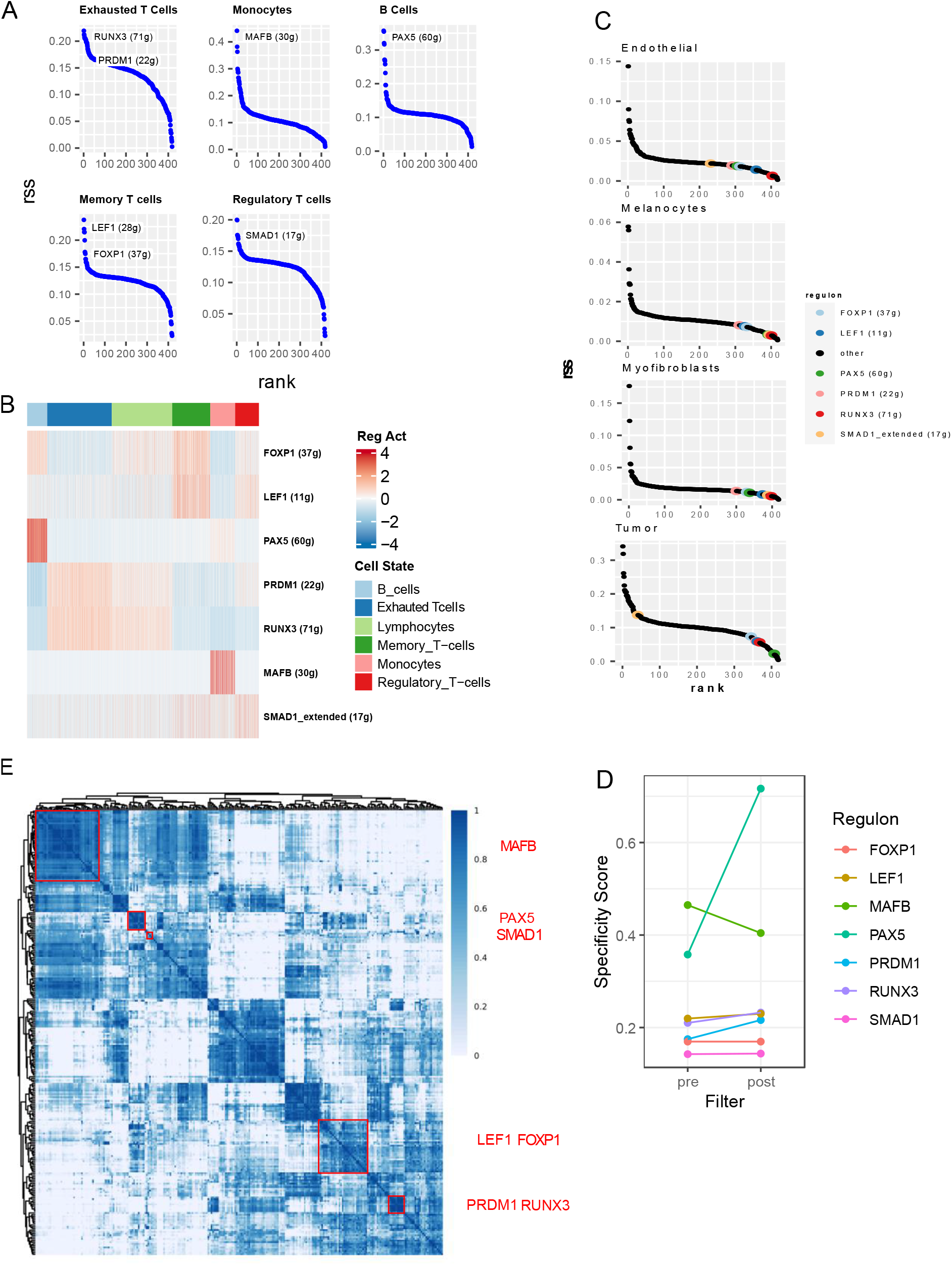
The regulons that define cell states of response. **A)** The rank of regulons in their associated cell state based on regulon specificity score (rss). The numbers in parenthesis denote the number of genes in the regulon before pruning. **B)** Relative activity level of each regulon across each of the DAseq-identified cell states. **C)** The rank of regulons in non-immune cells from an external BCC ICI dataset based on rss **D)** RSS for each regulon pre- and post-filtering of target genes according to gene markers of their associated cell type. **E)** Identified regulon modules based on regulon connection specificity index (CSI) matrix, along with the representative TFs. The color scale bar denotes the CSI score.

To distinguish between different immune cells and their functional states, the respective regulons must be specific to immune cells and not co-expressed in other cells present in the TME. We tested this in an independent ICI scRNA-seq dataset, containing basal cell carcinoma (BCC) samples with immune and non-immune cells.^10^ The specificity scores for each regulon were low in tumor cells, melanocytes, endothelial and myofibroblasts, suggesting that the presence of non-immune cells has little impact on the regulon specificity scores (Fig 2C).

Regulons inferred using regression are context-dependent and at risk of over-fitting the data. Complex regulons with a large number of genes are more likely to suffer from this phenomenon. To counteract this, we reduced the size of each regulon. Briefly, a the target genes of a TF were removed from a regulon if they were not upregulated in the predominant immune cell phenotypes of its associated cell state. As an example, PAX5 target genes were filtered according to genes upregulated in B cells. Using this approach, the complexity of each regulon was reduced, with an overall increase in specificity across all regulons (Fig 2D; Supplementary Table1).

To ensure that the regulons defining cell states quantified an independent signal, a pairwise similarity comparison of regulon activity across each cell was performed using the CSI (see Methods).^20^ The selected regulons clustered into distinct modules, suggesting that redundant regulons were not present (Fig 2E).

### Regulon Validation in Bulk RNA-seq datasets

To determine if these regulons could characterize response to ICIs in melanoma, we collected four datasets that contained pre-treatment transcriptomic data and therapy response information (n=209).^11–14^ The distribution of patients who had received mono- or combination therapy in the validation dataset (80% anti-PD1 or anti-CTLA4, 20% anti-CTLA4+PD-1) matched the discovery, scRNA-seq dataset (78% anti-PD1 or anti-CTLA4, 22% anti-CTLA4+PD-1) (Supplementary Table1).

The relative activity of each regulon was scored in each dataset independently using ssGSEA.^24^ Median centering and scaling of regulon scores were performed to reduce batch effects between datasets (Fig 3A), which were then combined for hierarchical clustering.

**Figure 3:**
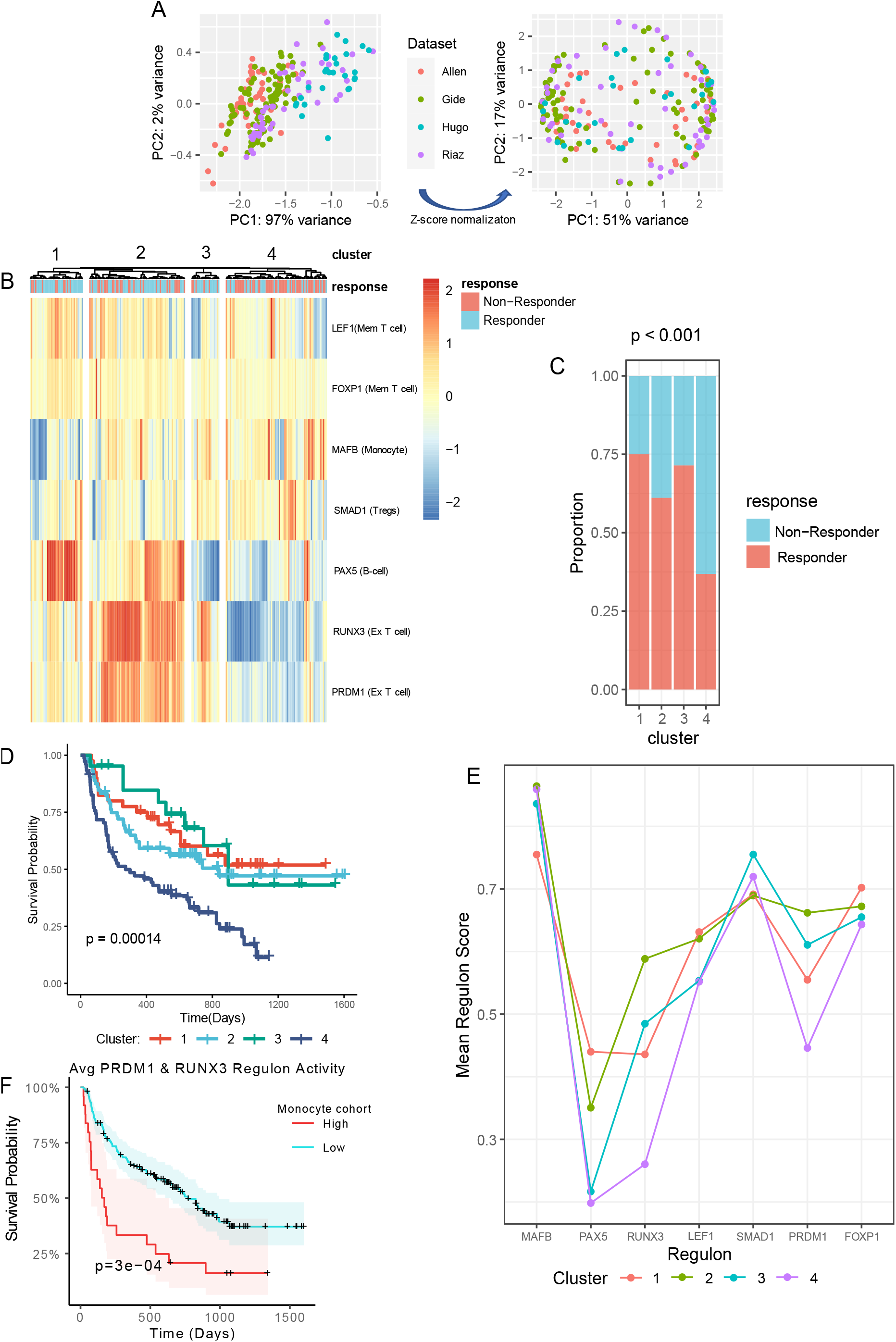
The regulons correlate with response in bulk RNA-seq samples. **A)** Principal component analysis (PCA) and clustering of patients from four datasets according to their regulon scores using ssGSEA, before (left image) and after (right image) z-score normalization. **B)** Hierarchical clustering of patients according to their ssGSEA inferred regulon scores, which are represented by the color scale bar. Pearson correlation was used as the distance metric. **C)** The proportion of responders and non-responders in each cluster. The p-value was calculated with a chi-squared test. **D)** Kaplan Meier survival curves for each cluster, compared using a log rank test. **E)** Mean regulon scores for each cluster identified using hierarchical clustering. **F)** Cox regression analysis comparing the association of PRDM1 and RUNX3 mean activity and survival in patients with high and low activity scores for MLCs. Statistical significance was inferred using the log rank test.

Four clusters with significantly different response outcomes (chi-squared test, p < 0.001) were identified (Fig 3B, 3C). The majority of patients in clusters 1,2 and 3 were responders (75%, 61% and 71%, respectively), whilst cluster 4 contained primarily non-responders (63%; Fig 3B, 3C). As expected, overall survival (OS) following ICI treatment was significantly improved in clusters containing primarily responders (Fig 3D).

Regulon scores were low in cluster 4, suggesting an immune desert phenotype, in which all compartments of the tumor are sparse for immune cells (Fig 3B, 3E).^40^ Cluster 1 contained high levels of productive memory T cell and B-cell states, with concomitant low values for immunosuppressive cell states (MLCs and Tregs), likely representing the infiltrated-inflamed phenotype (Fig 3B, 3E).^41^ Clusters 2 and 3 differed from cluster 1 in the number of MLCs and exhausted cell states present. We explored the relationship between these two cell states, and found by Cox-regression analysis that the exhausted T cell state predicts prognosis as a function of MLCs: higher numbers of MLCs are associated with worse survival (Fig 3F). This may explain why, despite having similar levels of other beneficial cell states, cluster 3, and to a greater extent cluster 2, had lower proportions of responders compared to cluster 1 (Fig 3C). As such, clusters 2 and 3 are likely the immune-inflamed phenotype, but the higher

### Monocyte Lineage cells and Exhausted T cells: Relationship Dynamics

We further investigated why PRDM1 and RUNX3 were associated with worse prognosis when MLCs were abundant. Patients from the scRNA-seq dataset were split into high and low cohorts according to the median number of cells in the exhausted T-cell state. The high cohort had significantly more MLCs (Fig 4A), suggesting that the exhausted cell state may be linked to the activity of MLCs.

**Figure 4:**
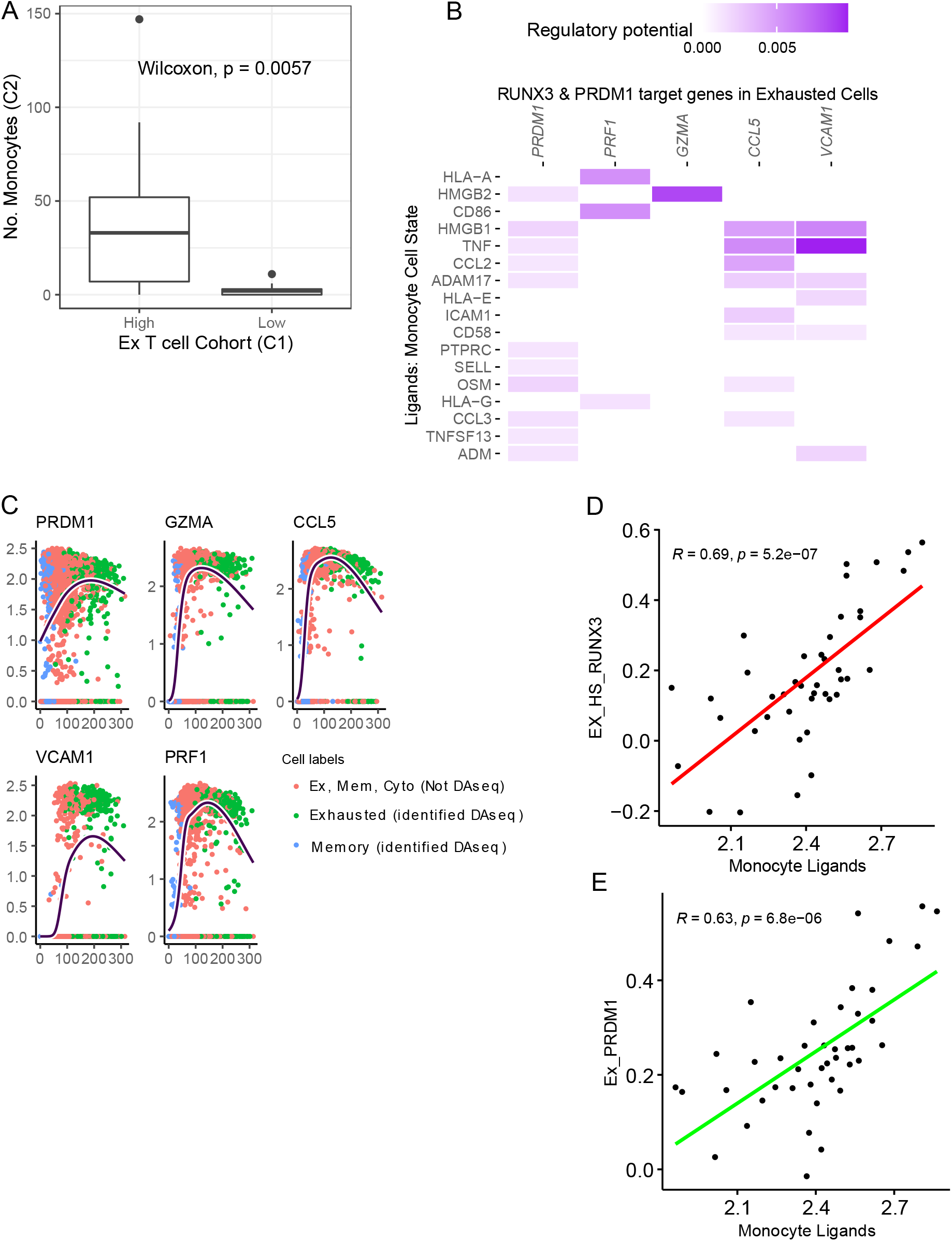
The relationship between MLCs and exhausted T cells. **A)** The number of MLCs in patients with either high or low exhausted T cell levels, compared using a Wilcoxon test. **B)** NicheNet inferred interactions between MLC genes and the PRDM1 and RUNX3 regulons in exhausted T cells. **C)** The average expression pattern of genes identified by NicheNet as regulated by MLCs across pseudo-time. The line represents the genes average expression with respect to pseudo-time, inferred using a generative additive model. The cells are annotated as Memory, Exhausted or Cyto (cytotoxic) T cells, and whether they were identified by DAseq. **D, E)** Pearson’s correlation between the top 5 MLC ligands from NicheNet, scored in the Van Allen dataset using ssGSEA, and the activity of RUNX3 and PRDM1.

An intercellular communication analysis, using NicheNet,^25^ identified genes whose expression in MLCs is associated with the expression of PRDM1 and RUNX3 gene targets in the exhausted T cell state. *HLA-A, HMGB1/2, CD86*, and *TNF* were the top scoring genes in MLCs and interacted with cytotoxic-related genes, such as *PRF1, GZMA* and *CCL5* (Fig 4B). We performed trajectory inference on memory, cytotoxic, and exhausted T cells and modelled the expression dynamics of cytotoxic-related genes along a dynamic biological timeline. The expression levels of *GZMA, PRDM1, CCL5* and *VCAM1* were low at early time-points, increased to a maximum, and then declined. Interestingly, exhausted T cells were the most prominent cell-type beyond the maximum expression point – when gene expression is decreasing (Fig 4C). This suggests that MLCs are a causative agent in an “activation-dependent exhaustion program” such as those reported previously.^42,43^

The Van Allen dataset was used to confirm the relationship between *HLA-A, HMGB2, CD86, HMGB1*, and *TNF* with PRDM1 and RUNX3 regulon activity: an ssGSEA inferred score for the MLC genes was strongly correlated with both PRDM1 and RUNX3 activity (R=0.63, R=0.69, respectively) (Fig 4D, 4E). Immune-related genes are likely to be correlated if they quantify the number of immune cells present in the TME. We accounted for this by fitting a multiple linear regression: the relationship of RUNX3 and PRDM1 activity with the MLC score was adjusted for a CIBERSORT inferred “Absolute Score”, which measures the total immune component present in a tumor. Importantly, the MLC score remained a significant predictor of RUNX3 and PRDM1 activities (*P*<.001), suggesting that this relationship is not confounded by the number of immune cells in the TME.

## DISCUSSION

Building on prior single-cell transcriptomic efforts to understand the relationship between the TME and ICI response,^9,43^ we comprehensively characterized the immune compartment in patients with metastatic melanoma prior to first-line ICI treatment. Unlike previous approaches, and in response to the inherent noise of scRNA-seq data, we used regulons to guide the characterization of cellular states. Our results showed that regulons constitute robust guides of cellular identity and have comparable performance to gene expression data in delineating immune cell phenotypes. Importantly, as the scoring of regulons can overcome batch and technical effects,^15^ our approach is likely better suited for validating findings from single cell analyses in a bulk RNA-seq context.

We identified four distinct immune cell states that were enriched in responder or non-responder tumor samples. As the cell states were identified using DAseq, they were not constrained to any predefined clusters, and thus, were not identified previously by Sade-Feldman et al.^9^ The exhausted CD8+ T cell state was enriched in non-responders, and was defined by RUNX3 and PRDM1, both of which demonstrate increased chromatin accessibility and expression in terminally exhausted cells.^5^ Terminally exhausted cells are the progeny of polyfunctional “progenitor exhausted” cells, and, in line with our findings, are considered unresponsive to ICIs.^4,5^

The exhausted CD8+ T cell state co-occurred with the MLC state. In melanoma, tumor-resident macrophages correlate with poor anti-PD1 response,^12,44^ whereas peripheral myeloid-derived suppressor cells correlate with poor anti-CTLA4 response.^9,45^ The mechanisms driving these associations are only partially understood. We report that the MLC state likely induces terminal exhaustion in CD8+ T cells. MLCs interact with the exhaustion programs in CD8+ T cells through ligands involved in antigen presentation (HLA-A), chronic inflammation (TNF) and negative co-stimulation (HMGB1 – a TIM3 ligand), all of which, are implicated in the development of T cell exhaustion.^46^ The expression of cytotoxic (*GZMA, VCAM1*) and activation (*CCL5*) genes across pseudo-time that interact with MLC ligands are in decline in the exhausted CD8+ T cell state. This decline is preceded by a maximum expression value, suggesting that these cells may be stimulated until they become desensitized to the co-stimulatory pathway signal. In the context of ICI response, we found that exhausted CD8+ T cells were associated with worse outcomes if high MLC levels were present, suggesting that the association between exhausted T cells and ICI resistance is inextricably linked to MLCs.

Two cell states were associated with a positive response to ICI: a B cell and memory T cell state. The memory cell state is defined by the LEF1 and FOXP1 regulons – TFs associated with early differentiated memory T cells. Upon mapping the organizational structure of CD8+ memory T cells, a memory cell state characterized by LEF1 and FOXP1 was identified.^47^ This cell state is described as potentially facilitating immunotherapy response through its self-renewal capabilities and propensity for activation and effector differentiation. Although the memory cell state identified here may not be transcriptionally identical - a potential consequence of different cellular ecosystems used for identifying the cell state (normal physiology vs melanoma), it is apparent from multiple studies that early differentiated, stem-cell like memory cells are important for response to ICIs.^4^

Reports on the impact of B cells on ICI response were initially inconsistent. It is now apparent that B cells can either bolster or negate ICIs, depending on their phenotype.^48^ In response to autologous melanoma secretomes, B cells differentiate into a plasmablast-like B cell population with upregulation of PAX5.^49^ In concordance with our PAX5-defined B cell state, the frequency of plasmablast-like B cells predicts response and survival to ICI by increasing PD-1+ T cell activation.^49^ We hypothesize that the PAX5 TF regulates a differentiation program in B cells exposed to melanoma stimuli, that, in conjunction with an ICI, can drive tumor clearance.

We demonstrated that highly resolved cellular states can identify patients likely to respond to ICIs. One can envision two scenarios where identifying these cellular states in patient samples can have clinical implications: 1) selecting patients for ICI therapy based on the presence of cellular states amenable to ICI response. 2) drug combinations that can transition cells from a resistant cellular state to a productive cellular state. Although cell modification strategies are an active area of research, particularly in the context of macrophages,^50^ future studies are needed to explore the dynamics between cellular states and drug combinations.

## Supporting information

graphical abstract

## Ethics approval and consent to participate

Not applicable

## Availability of data and material

Data are available in public, open access repositories. All data relevant to the study are included in the article or uploaded as online supplemental information. The public datasets used and/or analyzed during the current study are available in the GEO or Zenodo databases (https://www.ncbi.nlm.nih.gov/geo/; https://zenodo.org/record/4661265).

## Competing Interests

SD, AG, VVA, TS, CB and IB are employees of AstraZeneca. The remaining authors declare no competing interests.

## Funding

This publication has emanated from research conducted with the financial support of Precision Oncology Ireland, a Consortium of 5 Irish Universities, 6 Irish Charities, and 7 Industry Partners, which is part-funded by the Science Foundation Ireland Strategic Partnership Programme, under Grant number [18/SPP/3522]. This project has received co-funding from the Astra Zeneca.

## Authors’ contributions

DE, VZ, WK DB, IB and CB contributed to the conception and design of the analysis; DE and VZ collected and analyzed the data with feedback from all authors. DE wrote the paper with input from all authors. All authors have read and agreed to the published version of the manuscript.

## Acknowledgements

The graphical abstract was created with BioRender.com.

## List of Abbreviations

(ICI): Immune checkpoint inhibitor
(regulons): TF-directed co-expression networks
(TME): Tumor microenvironment
(Tregs): Regulatory T cells
(TAMs): Tumor-associated macrophages
(scRNA-seq): Single cell RNA-seq
(TF): Transcription factor
(UMI): Unique molecular identifier counts
(SNN): Shared nearest neighbor
(DA): Differentially abundant
(CR): Complete response
(PR): Partial response
(SD): Stable disease
(PD): Progressive disease
(CSI): Connection Specificity Index
(ssGSEA): Single Sample Gene Set Enrichment Analysis
(CS): Cell State
(MLCs): Monocyte lineage cells
(BCC): Basal cell carcinoma
(OS): Overall survival

## Notes

https://www.ncbi.nlm.nih.gov/geo/

https://zenodo.org/record/4661265

### Competing Interest Statement

The authors have declared no competing interest.

