## Supplementary material for "Small gene networks can delineate immune cell states and characterize immunotherapy response in melanoma": graphical abstract

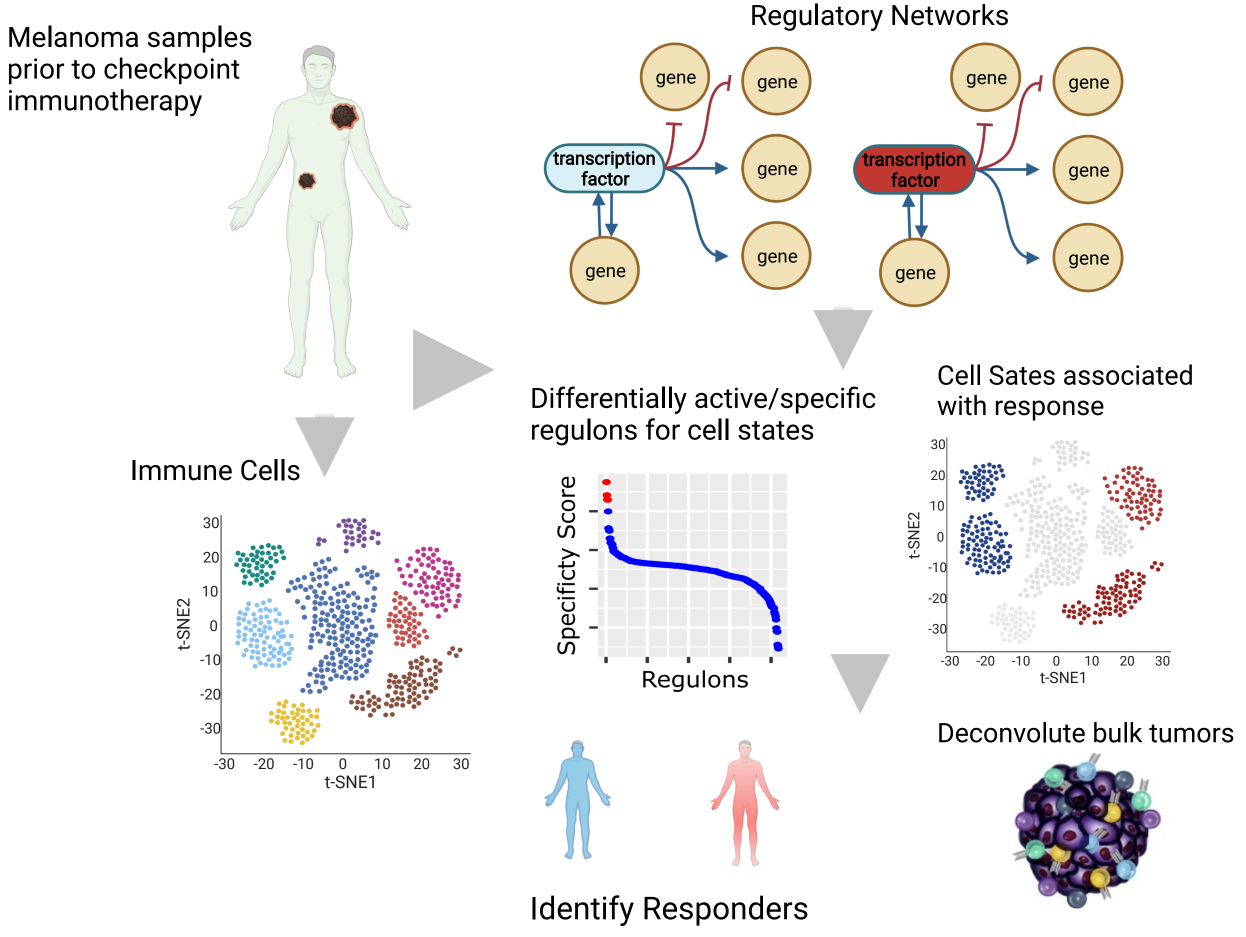


### Authors
Donagh Egan, Martina Kreileder, Myriam Nabhan, Luis F. Iglesias-Martinez, Simon Dovedi, Amit Grover, Viia Valge-Archer, Robert Wilkinson, Tim Slidel, Claus Bendtsen, Ian Barrett, Donal Brennan, Walter Kolch, and Vadim Zhernovkov

### Correspondence


### In Brief
Transcription Factor-directed gene networks can characterize immune cell states associated with immune checkpoint response, and be recovered in bulk RNAseq data to identify responders
